# Glycerol monolaurate ameliorated intestinal barrier and immunity in broilers by regulating intestinal inflammation, antioxidant balance, and intestinal microbiota

**DOI:** 10.1101/2021.05.19.444906

**Authors:** Linglian Kong, Zhenhua Wang, Chuanpi Xiao, Qidong Zhu, Zhigang Song

## Abstract

Extensive interactions occur between a poultry host and its gut microbiome. Glycerol monolaurate (GML) possesses a large range of antimicrobial and immunoregulatory properties. This study was conducted to investigate the impact of different doses of GML (basal diets complemented with 0, 300, 600, 900, or 1200 mg/kg GML) on growth performance, intestinal barrier, and cecal microbiota in broiler chicks. Results revealed that feed intake increased after 900 and 1200 mg/kg GML were administered during the entire 14-day experiment period. Dietary GML decreased crypt depth and increased the villus height-to-crypt depth ratio of the jejunum. In the serum and jejunum, supplementation with more than 600 mg/kg GML reduced interleukin-1β, tumor necrosis factor-α, and malondialdehyde levels and increased the levels of immunoglobulin G, jejunal mucin 2, total antioxidant capacity, and total superoxide dismutase. GML down-regulated jejunal interleukin-1β and interferon-*γ* expression and increased the mRNA level of zonula occludens 1 and occludin. A reduced expression of toll-like receptor 4 and a tendency of down-regulated nuclear factor kappa-B was shown in GML-treated groups. In addition, GML modulated the composition of the cecal microbiota of the broilers, improved microbial diversity, and increased the abundance of butyrate-producing bacteria. Spearman’s correlation analysis revealed that the genera *Barnesiella, Coprobacter, Lachnospiraceae, Faecalibacterium, Bacteroides, Odoriacter*, and *Parabacteroides* were related to inflammation and intestinal integrity. In conclusion, GML ameliorated intestinal morphology and barrier function in broiler chicks probably by regulating intestinal immune and antioxidant balance, as well as intestinal microbiota.

**IMPORTANCE:** Antibiotic residues and resistance issues led to the ban of antibiotic growth promoters. GML is considered an efficacious antibiotic growth promoter alternative for animal health and has the potential to become a unique fungicide owing to its established safety, antibacterial properties, and immunomodulatory capacity. Despite the potential of GML as an additive in poultry feed, little is known about the influence of GML on cecal microbiota in broilers. The significance of our research was to determine the microbial mechanism by which GML worked.

## INTRODUCTION

Antibiotics play a significant role in disease prevention and growth promotion in the poultry industry. Despite increasing demand for poultry, antibiotic residues and resistance issues led to the ban of antibiotic growth promoters (1), pressuring the industry to find alternatives for maintaining poultry flock health. One promising approach is immune modulation, in which natural host mechanisms are exploited to enhance and modulate bird’s immune response (2). Targeted dietary supplementation or using a feed additive may be useful in the immunomodulation of the immune system. These ingredients can reduce the negative impacts of environmental stressors on animal immune systems and production performance (3). For instance, antimicrobial peptides are feed additives that neutralize lipopolysaccharide (LPS) from *Pseudomonas aeruginosa* at the cellular level and significantly inhibit tumor necrosis factor (TNF-α) and nitric oxide (NO) production in the macrophages of LPS-treated mice (4). Extensive research has been carried out to evaluate an array of products as alternatives to antibiotic growth promoters; such products, including food industry by-products, plant metabolites, non-digestible oligosaccharides, natural by-products, essential minerals, amino acids, medicinal herbs, organic acids, and essential oils, can at least partially alter immune function in poultry (2).

Glycerol monolaurate (GML), a fatty acid composed of glycerol and lauric acid, possesses a large range of antimicrobial and immunoregulatory properties (5). Glycerol monolaurate is considered a food-safe emulsifier endorsed by the Food and Drug Administration and recognized as a nontoxic compound even at relatively high dose levels (6). Recent studies revealed that GML is an efficacious antibiotic growth promoter alternative for animal health (7) and has the potential to become a unique fungicide owing to its established safety, antibacterial properties, and immunomodulatory capacity (8). Glycerol monolaurate with a supplementary dose of up to 5 g/kg to basal diets can enhance the immune status and intestinal histomorphology of broilers (3). In vivo, growth performance and intestinal development are improved by dietary GML in mice and laying hens through intestinal microbiota alteration (6).

Despite the potential of GML as an additive in poultry feed (9), little is known on the influence of GML on intestinal barrier and cecal microbiota in broilers. Extensive interactions occur between a poultry host and its gut microbiome, particularly during exchange of nutrients and modulation of host gut morphology, physiology, and immunity (10). We hypothesized that GML can improve immunity, growth performance, and health status by altering gut microbiota. Therefore, the present study was designed to investigate the effects of different doses of GML on performance, immunity, and antioxidant capacity, as well as intestinal barrier and microbiota in broilers.

## RESULTS

### Growth performance

As shown in Table 3, FI rates increased in the 600, 900, and 1200 mg/kg GML-treated groups compared with those in the 300 mg/kg group (*P* < 0.05) after 7–14 days of treatment but showed no significant difference from those of the control group (*P* > 0.05). However, the administration of 900 and 1200 mg/kg GML increased FI compared to the control and 300 mg/kg GML-treated group during the overall period (*P* < 0.05). Dietary GML did not affect the BW, BWG, and FCR of broiler chicks. (*P* > 0.05).

**TABLE 1.** Ingredient composition and nutritional components of the basal diet

| Items | Content (%) |
| --- | --- |
| Ingredient |  |
| Corn | 60 |
| Soybean meal (43) | 28.8 |
| Corn protein flour (60) | 5.3 |
| Salt | 0.16 |
| Baking soda | 0.2 |
| Limestone | 1.3 |
| Dicalcium phosphate | 0.75 |
| Soybean oil | 2.2 |
| Vitamin premix | 0.03 |
| Mineral premix | 0.2 |
| Choline chloride (50%) | 0.1 |
| Methionine | 0.23 |
| Lysine (70%) | 0.58 |
| Threonine (98.5%) | 0.134 |
| Phytase (20000U) | 0.02 |
| Total | 100 |
| Nutritional level |  |
| Metabolizable energy | 2850 (kcal/kg) |
| Crude protein | 21.5 |
| Lysine | 1.26 |
| Methionine | 0.57 |
| Calcium | 0.60 |
| Total phosphorus | 0.60 |
| Available phosphorus | 0.40 |

Provided for per kilogram of compound diet: vitamin A, 12000 IU; vitamin D3, 5000 IU; vitamin E, 80 mg; VK, 3.2 mg; vitamin B1, 3.2 mg; vitamin B2, 8.6 mg; nicotinic acid, 65 mg; pantothenic acid, 20 mg; vitamin B6, 4.3 mg; biotin, 0.22 mg; folic acid, 2.2 mg; vitamin B12, 0.017 mg; I, 1.25 mg; Fe, 20 mg; Mn, 120 mg; Se, 0.3 mg ; Zn, 110 mg. Nutrition level was the calculated value.

**TABLE 2.**
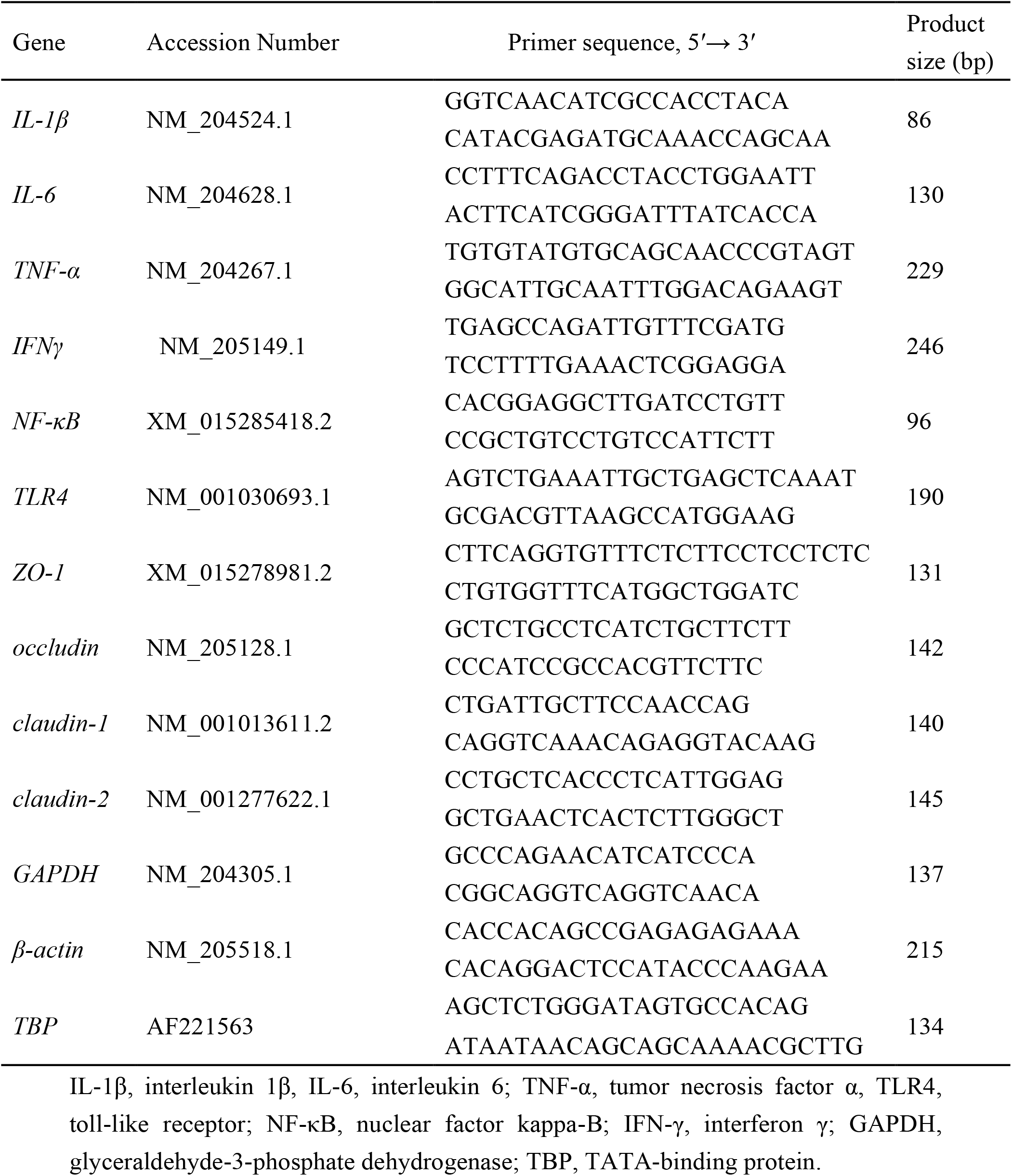
Nucleotide sequences for real-time PCR primers

**TABLE 3.** Effects of dietary treatment on growth performance in 7- and 14-day-old broilers

| Items | Treatments |  |  |  |  | *P*-value |
| --- | --- | --- | --- | --- | --- | --- |
|  | Control | 300 mg/kg | 600 mg/kg | 900 mg/kg | 1200 mg/kg |  |
| BW (g/bird) |  |  |  |  |  |  |
| 1d | 45.71±0.27 | 45.65±0.15 | 45.68±0.08 | 45.67±0.19 | 45.67±0.26 | 0.992 |
| 7d | 154.82±6.25 | 151.19±6.66 | 148.71±4.84 | 150±8.19 | 152.99±2.22 | 0.437 |
| 14d | 401.42±8.99 | 389.23±8.77 | 392.67±16.09 | 392.54±7.82 | 397.07±16.78 | 0.149 |
| FI (g/bird) |  |  |  |  |  |  |
| 1 to 7d | 145.74±7.85 | 148.6±2.78 | 143.01±6.03 | 150.34±1.05 | 155.88±5.04 | 0.123 |
| 7 to 14d | 354.74±13.39<sup>ab</sup> | 337.05±18.91<sup>b</sup> | 367.62±4.35<sup>a</sup> | 375.46±13.88<sup>a</sup> | 366.22±27.4<sup>a</sup> | 0.013 |
| 1 to 14d | 500.04±13.3<sup>bc</sup> | 485.75±17.41<sup>c</sup> | 511.82±17.56<sup>ab</sup> | 525.64±6.91<sup>a</sup> | 521.54±14.46<sup>a</sup> | 0.001 |
| BWG (g/bird) |  |  |  |  |  |  |
| 1 to 7d | 116.11±6.15 | 120.54±6.75 | 118.05±4.87 | 119.33±8.06 | 122.32±2.34 | 0.44 |
| 7 to 14d | 237.6±9.36 | 221.38±8.04 | 228.95±12.93 | 227.54±4.09 | 229.08±15.14 | 0.202 |
| 1 to 14d | 356.71±9.07 | 343.6±8.91 | 347±16.11 | 346.88±7.66 | 351.4±16.93 | 0.155 |
| FCR (g/g) |  |  |  |  |  |  |
| 1 to 7d | 1.25±0.04 | 1.24±0.06 | 1.21±0.02 | 1.26±0.1 | 1.27±0.04 | 0.343 |
| 7 to 14d | 1.49±0.07 | 1.56±0.1 | 1.62±0.03 | 1.62±0.09 | 1.56±0.12 | 0.18 |
| 1 to 14d | 1.41±0.04 | 1.41±0.07 | 1.47±0.03 | 1.50±0.01 | 1.48±0.07 | 0.13 |

BW, body weight; FI, feed intake; FCR, feed conversion ratio.
Values were expressed as mean ± SD (n = 6).
<sup>a,b,c</sup> Different superscripts in the same line indicate significant differences (*P* < 0.05).

### Intestinal morphology analysis

As shown in Table 4, GML decreased CD (*P* < 0.05) and increased VCR (*P* < 0.05) in the jejunum on day 7 and 14 with increasing dose (600, 900, and 1200 mg/kg). No effect was observed on jejunal VH after dietary treatment with GML (*P* > 0.05).

**TABLE 4.** Effects of dietary treatment on the jejunal morphology of 7- and 14-day-old broilers

| Items | Treatments |  |  |  |  | *P*-value |
| --- | --- | --- | --- | --- | --- | --- |
|  | CON | 300 mg/kg | 600 mg/kg | 900 mg/kg | 1200 mg/kg |  |
| 7 d of age |  |  |  |  |  |  |
| VH (μm) | 659.86±96.54 | 708.71±64.12 | 706.49±84.1 | 684.76±77.11 | 675.15±76.88 | 0.824 |
| CD (μm) | 107.01±17.21<sup>a</sup> | 101.01±15.22<sup>ab</sup> | 83.39±16.79<sup>bc</sup> | 67.32±11.39<sup>cd</sup> | 60.48±9.82<sup>d</sup> | <0.001 |
| VCR | 6.24±1.02<sup>c</sup> | 7.11±0.95<sup>bc</sup> | 8.66±1.45<sup>b</sup> | 10.34±1.62<sup>a</sup> | 11.3±1.51<sup>a</sup> | <0.001 |
| 14 d of age |  |  |  |  |  |  |
| VH (μm) | 745.08±71.15 | 697.3±86.11 | 670.03±99.88 | 686.95±28.38 | 708.35±61.9 | 0.765 |
| CD (μm) | 102.33±12.34<sup>a</sup> | 89.9±7.99<sup>b</sup> | 73.28±0.86<sup>c</sup> | 66.33±4.16<sup>c</sup> | 69.1±6.76<sup>c</sup> | <0.001 |
| VCR | 7.3±0.23<sup>b</sup> | 7.84±1.39<sup>b</sup> | 8.88±1.58<sup>ab</sup> | 10.37±0.47<sup>a</sup> | 10.34±1.44<sup>a</sup> | 0.011 |

VH, villus height; CD, crypt depth; VCR, villus height to crypt depth ratio.
Values were expressed as mean ± SD (n = 6).
<sup>a,b,c,d</sup> Different superscripts in the same line indicate significant differences (*P* < 0.05).

### Serum and intestinal biochemical indicators

On days 7 and 14, dietary treatment with 600, 900, and 1200 mg/kg GML reduced serum IL-1β level (*P* < 0.05) compared with the level in the control group (Table 5). A reduction (*P* < 0.05) of serum TNF-α level was recorded in 7-day-old broilers treated with 900 and 1200 mg/kg GML. In addition, 1200 mg/kg GML increased serum IgG and jejunal mucin 2 levels relative to those of the control on day 14.

**TABLE 5.** Effects of dietary treatment on biochemical indicators in the sera and jejuna of 7- and 14-day-old broilers

| Items | Treatments |  |  |  |  | *P*-value |
| --- | --- | --- | --- | --- | --- | --- |
|  | CON | 300mg/kg | 600mg/kg | 900mg/kg | 1200mg/kg |  |
| 7 d of age (ng/mL) |  |  |  |  |  |  |
| IL-1β | 597.46±33.97<sup>a</sup> | 559.36±45.06<sup>ab</sup> | 531.91±34.96<sup>b</sup> | 541.96±37.13<sup>b</sup> | 525.78±52.65<sup>b</sup> | 0.028 |
| IL-6 | 29.05±2.08 | 28.9±1.71 | 28.62±1.67 | 29.73±1.1 | 29.57±1.05 | 0.897 |
| TNF-α | 81.36±6.26<sup>a</sup> | 79.03±3.78<sup>ab</sup> | 78.27±2.9<sup>ab</sup> | 75.44±7.37<sup>bc</sup> | 72.75±3.49<sup>c</sup> | 0.029 |
| IgG | 1934.38±153.59 | 1911.25±107 | 1832.5±175.64 | 1953.13±157.39 | 1829.38±160.03 | 0.35 |
| sIgA | 26.01±3.4 | 26.48±3.09 | 25.7±2.6 | 26.6±2.61 | 25.32±2.82 | 0.891 |
| Muc2 | 213.65±8.99 | 209.97±8.15 | 201.65±13.8 | 202.8±17.57 | 197.98±13.87 | 0.219 |
| 14 d of age (ng/mL) |  |  |  |  |  |  |
| IL-1β | 629.71±27.08<sup>a</sup> | 583.17±35.5<sup>ab</sup> | 566.84±39.61<sup>b</sup> | 543.68±56.89<sup>b</sup> | 560.59±51.03<sup>b</sup> | 0.018 |
| IL-6 | 29.49±2.86 | 29.63±0.68 | 29.63±1.22 | 30.96±1.94 | 30.2±1.23 | 0.746 |
| TNF-α | 76.9±8.37 | 73.69±3.46 | 78.36±1.64 | 77.96±2.12 | 78.21±3.92 | 0.162 |
| IgG | 1898.13±195.47<sup>b</sup> | 1813.13±145.67<sup>b</sup> | 1941.88±167.71<sup>ab</sup> | 1939.38±202.95<sup>ab</sup> | 2115±212.94<sup>a</sup> | 0.042 |
| sIgA | 24.56±1.85 | 25.1±2.53 | 26.91±1.38 | 25.81±2.13 | 25.73±1.93 | 0.268 |
| Muc 2 | 196.06±8.96<sup>b</sup> | 192.53±12.58<sup>b</sup> | 202.68±15.33<sup>ab</sup> | 201.31±12.18<sup>ab</sup> | 212.2±10.18<sup>a</sup> | 0.028 |

IL-1β , interleukin 1 $\beta$ ; TNF-α , tumor necrosis factor $\alpha$ ; IgG, immunoglobulin G; sIgA, secreted immunoglobulin A; Muc 2, mucin 2.
Values were expressed as mean ± SD (n = 12).
<sup>a,b,c</sup> Different superscripts in the same line indicate significant differences ( $P < 0.05$ ).

### Antioxidant capacity in serum and jejunum

As shown in Table 6, dietary GML reduced MDA content in the serum and jejunum on 14-day-old broilers (*P* < 0.05). Serum SOD level increased in broilers fed with 1200 mg/kg GML on day 7 (*P* < 0.05) relative to that of the control and tended to improve 14 days after GML supplementation (*P* < 0.1). In the jejuna of 7-day-old broilers, dietary treatment with GML increased SOD levels (*P* < 0.05), and the highest level was found in the 1200 mg/kg group (*P* < 0.05). A higher T-AOC level was observed in the jejuna of the 600 and 1200 mg/kg groups on day 7 (*P* < 0.05) and in the sera after 900 and 1200 mg/kg GML addition on day 14 (*P* < 0.05).

**TABLE 6.** Effects of dietary treatment on antioxidant capacity in 7- and 14-day-old broilers

| Items | Treatments |  |  |  |  | *P*-value |
| --- | --- | --- | --- | --- | --- | --- |
|  | CON | 300 mg/kg | 600 mg/kg | 900 mg/kg | 1200 mg/kg |  |
| Serum, 7 d of age |  |  |  |  |  |  |
| MDA (nmol/ml) | 5.88±1.87 | 5.29±0.68 | 5.99±1.36 | 5.67±0.43 | 5.08±0.68 | 0.303 |
| SOD (U/ml) | 67.41±14.18<sup>b</sup> | 77.19±15.5<sup>ab</sup> | 65.4±4.71<sup>b</sup> | 82.87±11.1<sup>ab</sup> | 93.84±17.69<sup>a</sup> | 0.039 |
| T-AOC (mmol/ml) | 0.56±0.06 | 0.54±0.11 | 0.52±0.08 | 0.58±0.1 | 0.63±0.11 | 0.269 |
| 14 d of age |  |  |  |  |  |  |
| MDA (nmol/ml) | 6.6±1.39<sup>a</sup> | 4.64±1.23<sup>b</sup> | 4.07±0.72<sup>b</sup> | 4.51±0.56<sup>b</sup> | 4.95±1.12<sup>b</sup> | 0.005 |
| SOD (U/ml) | 72.51±6.54 | 84.33±28.02 | 89.43±9.3 | 101.29±26.88 | 111±21.96 | 0.071 |
| T-AOC (mmol/ml) | 0.41±0.05<sup>b</sup> | 0.47±0.05<sup>ab</sup> | 0.49±0.11<sup>ab</sup> | 0.52±0.04<sup>a</sup> | 0.5±0.04<sup>a</sup> | 0.005 |
| Jejunum, 7 d of age |  |  |  |  |  |  |
| MDA (nmol/mgprot) | 0.36±0.17 | 0.65±0.36 | 0.84±0.67 | 0.69±0.47 | 0.65±0.28 | 0.170 |
| SOD (U/mgprot) | 2.47±0.48<sup>d</sup> | 3.35±0.55<sup>c</sup> | 4.11±0.95<sup>ab</sup> | 3.6±0.59<sup>bc</sup> | 4.49±0.84<sup>a</sup> | <0.001 |
| T-AOC (mmol/mgprot) | 0.08±0.01<sup>b</sup> | 0.1±0.02<sup>b</sup> | 0.12±0.03<sup>a</sup> | 0.1±0.02<sup>b</sup> | 0.16±0.08<sup>a</sup> | 0.029 |
| 14 d of age |  |  |  |  |  |  |
| MDA (nmol/mgprot) | 0.98±0.21<sup>a</sup> | 0.6±0.08<sup>b</sup> | 0.56±0.16<sup>b</sup> | 0.55±0.16<sup>b</sup> | 0.6±0.14<sup>b</sup> | <0.001 |
| SOD (U/mgprot) | 3.72±0.65 | 3.77±0.7 | 3.76±0.9 | 4.01±0.74 | 3.39±0.55 | 0.556 |
| T-AOC (mmol/mgprot) | 0.08±0.01 | 0.08±0.01 | 0.08±0.02 | 0.08±0.02 | 0.07±0.01 | 0.686 |

MDA, malondialdehyde; SOD, superoxide dismutase; T-AOC, total antioxidant capacity; mgprot, mg of protein.
Values were expressed as mean ± SD (n = 12).
<sup>a,b,c,d</sup> Different superscripts in the same line indicate significant differences (*P* < 0.05).

### Relative mRNA expression of jejunal genes

On 7-day-old broilers, dietary treatment with 1200 mg/kg GML down-regulated (*P* < 0.05) jejunal *IL-1β* and *interferon (IFN-γ)* expression compared with that in the control (Table 7). Moreover, 600 mg/kg GML-treated broilers showed higher *zonula occludens (ZO)-1* and *occludin* expression levels than the control (*P* < 0.05). The mRNA level of *occludin* increased in the jejunum with 1200 mg/kg GML supplementation (*P* < 0.05). On 14-day-old broilers, *IL-1β* expression decreased in the 1200 mg/kg group (*P* < 0.05), and *IFN-γ* expression decreased in the 600, 900, and 1200 mg/kg groups (*P* < 0.05). Jejunal *occludin* expression was not altered by GML supplementation, compared with that in the control (*P* > 0.05), but difference in jejunal *occludin* expression between GML-treated groups was observed (*P* < 0.05). The expression of toll-like receptor4 (TLR4) was decreased in 600 and 1200 mg/kg GML-treated broilers (*P* < 0.05). In addition, dietary GML tended to reduce the expression of jejunal nuclear factor kappa-B (NF-κB) in 14-day-old broilers (*P* < 0.1).

**TABLE 7.** Effects of dietary treatment on jejunal gene expression of 7- and 14-day-old broilers

| Item | Treatment |  |  |  |  | *P*-value |
| --- | --- | --- | --- | --- | --- | --- |
|  | CON | 300 mg/kg | 600 mg/kg | 900 mg/kg | 1200 mg/kg |  |
| 7 d of age |  |  |  |  |  |  |
| *IL-1β* | 1±0.19<sup>a</sup> | 0.77±0.23<sup>ab</sup> | 0.96±0.25<sup>a</sup> | 0.8±0.34<sup>ab</sup> | 0.55±0.07<sup>b</sup> | 0.019 |
| *IL-6* | 1±0.24 | 1.49±0.23 | 1.55±0.35 | 1.32±0.16 | 1.16±0.27 | 0.281 |
| *TNF-α* | 1±0.29 | 1.01±0.23 | 0.92±0.27 | 0.76±0.06 | 0.72±0.05 | 0.256 |
| *IFN-γ* | 1±0.31<sup>a</sup> | 0.96±0.57<sup>ab</sup> | 1.28±0.33<sup>a</sup> | 0.85±0.32<sup>ab</sup> | 0.59±0.19<sup>b</sup> | 0.047 |
| *ZO-1* | 1±0.1<sup>b</sup> | 1.13±0.19<sup>ab</sup> | 1.31±0.17<sup>a</sup> | 1.17±0.21<sup>ab</sup> | 1.03±0.07<sup>ab</sup> | 0.029 |
| *occludin* | 1±0.18<sup>b</sup> | 1.54±0.12<sup>a</sup> | 1.31±0.09<sup>a</sup> | 1.04±0.11<sup>b</sup> | 1.31±0.32<sup>a</sup> | 0.002 |
| *claudin-1* | 1±0.13 | 0.99±0.18 | 1.29±0.34 | 1.02±0.35 | 1.35±0.66 | 0.374 |
| *claudin-2* | 1±0.29 | 0.96±0.32 | 1.22±0.48 | 1.22±0.43 | 1.17±0.28 | 0.689 |
| *TLR4* | 1±0.2 | 1.06±0.32 | 0.91±0.07 | 0.83±0.21 | 1.01±0.13 | 0.501 |
| *NF-κB* | 1±0.13 | 0.97±0.25 | 1.11±0.22 | 1.01±0.23 | 0.88±0.17 | 0.443 |
| 14 d of age |  |  |  |  |  |  |
| *IL-1β* | 1±0.55 | 0.62±0.18 | 0.75±0.29 | 0.79±0.27 | 0.93±0.21 | 0.208 |
| *IL-6* | 1±0.18<sup>a</sup> | 0.87±0.26<sup>ab</sup> | 0.84±0.17<sup>ab</sup> | 1.09±0.23<sup>a</sup> | 0.65±0.23<sup>b</sup> | 0.037 |
| *TNF-α* | 1±0.2 | 0.94±0.21 | 1.04±0.15 | 0.88±0.26 | 1.07±0.24 | 0.548 |
| *IFN-γ* | 1±0.51<sup>a</sup> | 0.61±0.19<sup>ab</sup> | 0.52±0.13<sup>b</sup> | 0.51±0.25<sup>b</sup> | 0.45±0.23<sup>b</sup> | 0.019 |
| *ZO-1* | 1±0.08 | 1.02±0.16 | 1.13±0.18 | 1.2±0.25 | 1.34±0.35 | 0.111 |
| *occludin* | 1±0.12<sup>abc</sup> | 0.86±0.19<sup>bc</sup> | 0.77±0.16<sup>c</sup> | 1.26±0.13<sup>a</sup> | 1.13±0.2<sup>ab</sup> | 0.001 |
| *claudin-1* | 1±0.37 | 0.92±0.25 | 0.88±0.18 | 1.09±0.36 | 0.84±0.21 | 0.494 |
| *claudin-2* | 1±0.16 | 0.99±0.15 | 1±0.26 | 1.24±0.39 | 1.24±0.44 | 0.419 |
| *TLR4* | 1±0.35<sup>a</sup> | 0.69±0.18<sup>ab</sup> | 0.63±0.15<sup>b</sup> | 0.82±0.14<sup>ab</sup> | 0.65±0.09<sup>b</sup> | 0.049 |
| *NF-κB* | 1±0.33 | 0.7±0.12 | 0.79±0.08 | 0.84±0.13 | 0.8±0.17 | 0.093 |

IL-1β, interleukin 1β, IL-6, interleukin 6; TNF-α, tumor necrosis factor α, TLR4, toll-like receptor; NF-κB, nuclear factor kappa-B; IFN-γ, interferon γ; GAPDH, glyceraldehyde-3-phosphate dehydrogenase; TBP, TATA-binding protein.
Values were expressed as mean ± SD (n = 6).
<sup>a,b,c</sup> Different superscripts in the same line indicate significant differences ( $P < 0.05$ ).

### Composition and community diversity of cecal microbiota

On 7-day-old broilers, 378 OTUs were found among the five groups, and 626, 682, 843, 818, and 827 specific OTUs were unique to the control, 300, 600, 900, and 1200 mg/kg GML-treated groups, respectively (Fig. 1A). On 14 days of age, 938, 845, 616, 964, and 973 specific OTUs existed respectively in the Con, 300, 600, 900, and 1200 mg/kg GML-treated groups (Fig. 1A). As shown in Table 8, diets supplemented with 600, 900, and 1200 mg/kg GML increased Chao 1 and observed indices in 7-day-old broilers compared with the control (*P* < 0.05). However, no significant differences between the control and GML-treated groups were observed for alpha diversity on 14-day-old broilers (*P* > 0.05). Broilers fed with 1200 mg/kg GML had the highest (*P* < 0.05) Chao 1, PD, observed, Shannon, and Simpson indices among the GML-treated groups. On day 7, unweighted UniFrac-based nonmetric multidimensional scaling analysis indicated no obvious difference (*P* > 0.05) in the β diversity of cecal microbiota between each group (Fig. 1B). In the 14-day-old broilers, the microbiomes in the 900 and 1200 mg/kg GML groups were completely separated from the control and 300 mg/kg on the NMDS2 axes (*P* = 0.003; Fig. 1B). PERMANOVA analysis based on unweighted UniFrac distance revealed that the cecal microbiota in the 600, 900, and 1200 mg/kg GML-treated groups had a higher β diversity index than that in the control on day 7 (*P* < 0.05) and in the 900 and 1200 mg/kg GML-treated groups on day 14 (*P* < 0.05; Fig. 1C).

**TABLE 8.** Effects of GML on cecal microbiota α-diversity of broilers

| Item | Treatment |  |  |  |  | *P*-value |
| --- | --- | --- | --- | --- | --- | --- |
|  | CON | 300mg/kg | 600mg/kg | 900mg/kg | 1200mg/kg |  |
| 7 d of age |  |  |  |  |  |  |
| Chao 1 | 410.11±33.93<sup>b</sup> | 435.31±78.4<sup>ab</sup> | 497.67±53.9<sup>a</sup> | 506.31±71.22<sup>a</sup> | 501.08±61.47<sup>a</sup> | 0.029 |
| Pd | 28.09±2.25<sup>ab</sup> | 27.39±4.35<sup>ab</sup> | 26.76±3.77<sup>b</sup> | 27.28±1.88<sup>b</sup> | 31.02±2.54<sup>a</sup> | 0.112 |
| Observed | 409.67±34<sup>b</sup> | 434.17±78.23<sup>ab</sup> | 496.33±54.08<sup>a</sup> | 505±70.25<sup>a</sup> | 499.5±60.82<sup>a</sup> | 0.029 |
| Shannon | 6.22±0.31 | 6.29±0.62 | 6.67±0.6 | 6.7±0.59 | 6.6±0.42 | 0.393 |
| Simpson | 0.96±0.01 | 0.96±0.03 | 0.97±0.02 | 0.97±0.02 | 0.97±0.01 | 0.649 |
| 14 d of age |  |  |  |  |  |  |
| Chao 1 | 548.04±45.24<sup>a</sup> | 550.24±83.38<sup>a</sup> | 452.16±48.86<sup>b</sup> | 548.93±109.31<sup>ab</sup> | 612.15±45.41<sup>a</sup> | 0.018 |
| Pd | 34.35±13.06<sup>ab</sup> | 32.23±8.92<sup>ab</sup> | 28.18±2.59<sup>b</sup> | 32.16±4.49<sup>ab</sup> | 35.03±4.08<sup>a</sup> | 0.111 |
| Observed | 547.33±45.56<sup>a</sup> | 548.67±83.56<sup>a</sup> | 451.33±48.63<sup>b</sup> | 547.33±109.79<sup>ab</sup> | 611±45.22<sup>a</sup> | 0.018 |
| Shannon | 5.36±0.57<sup>a</sup> | 5.03±1.01<sup>ab</sup> | 4.51±0.36<sup>b</sup> | 4.99±0.51<sup>ab</sup> | 5.57±0.46<sup>a</sup> | 0.033 |
| Simpson | 0.89±0.06<sup>ab</sup> | 0.83±0.13<sup>ab</sup> | 0.85±0.04<sup>b</sup> | 0.87±0.02<sup>b</sup> | 0.91±0.03<sup>a</sup> | 0.153 |

Values were expressed as mean ± SD (n = 6).
<sup>a,b</sup> Different superscripts in the same line indicate significant differences according to the Kruskal-Wallis test ( $P < 0.05$ ).

**FIG 1.**
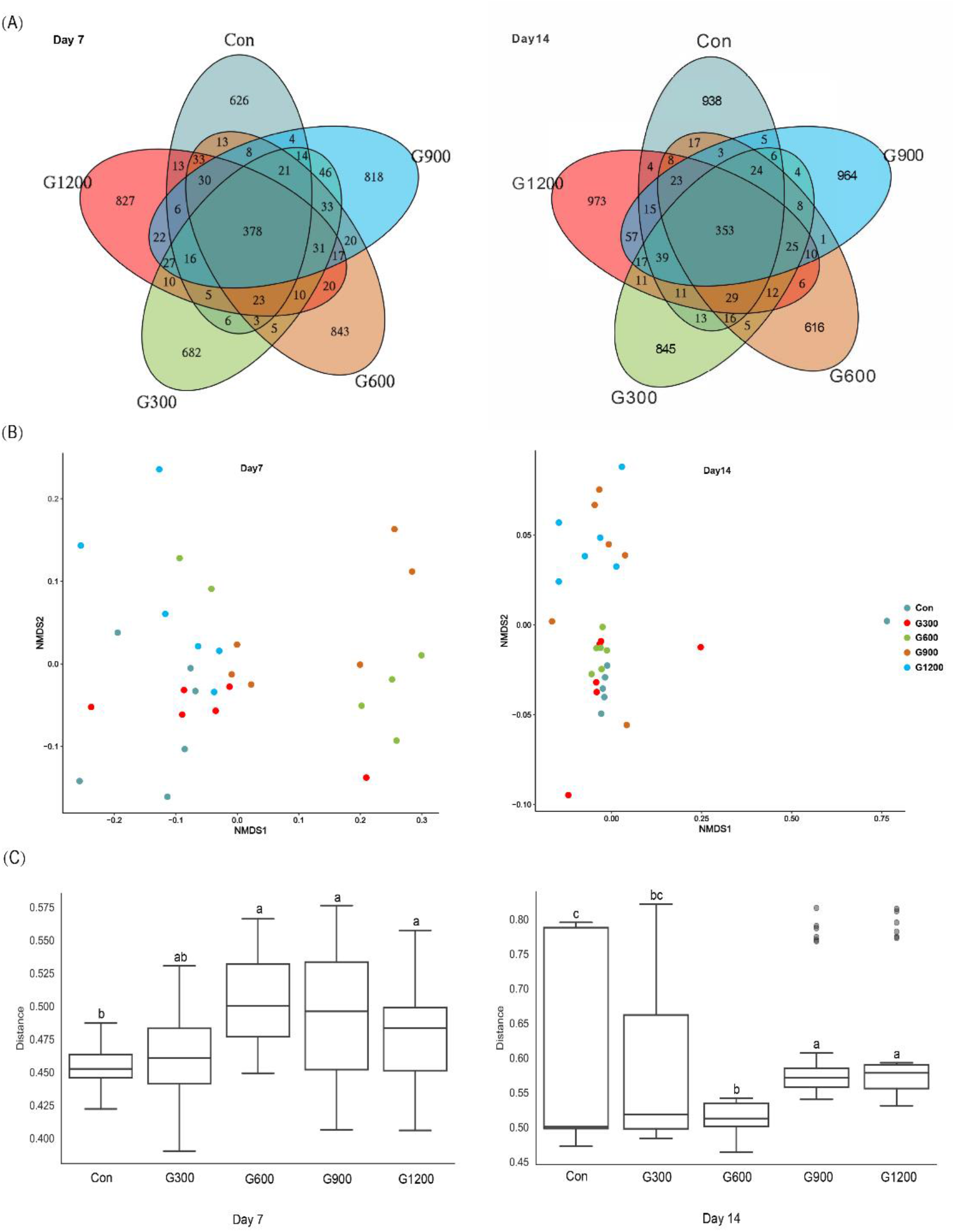
Dietary GML altered the composition and community diversity of cecal microbiota. (A) Venn diagram between treatments on OTUs level. (B) NMDS analysis based on unweighted Unifrac distance. (C) PERMANOVA analysis based on unweighted Unifrac distance. Con, basal diet; G300, 600, 900, and 1200, basal diets complemented with 300, 600, 900, or 1200 mg/kg GML.

The relative abundance of community at the phylum level is shown in Fig. 2A. At the phylum level, the *Firmicutes/Bacteroidetes* ratio decreased in the ceca of 7-day-old broilers after 600 and 1200 mg/kg GML supplementation (*P* < 0.05) but was not altered in each group on day 14 (*P* > 0.05; Fig. 2B). Supplementation with 600 and 1200 mg/kg GML increased the amount of *Bacteroidetes* relative to the amounts in the control, 300, and 900 mg/kg groups (*P* < 0.05; Fig. 2C). The relative abundance of *Actinobacteria* decreased in the ceca of the 14-day-old broilers that received 600 and 900 mg/kg GML (*P* < 0.05; Fig. 2C). At the genus level, the 300 mg/kg GML-treated group was enriched with *Barnesiella* and *CHKCI001* on days 7 and 14 (*P* < 0.05), respectively (Fig. 3B). The proportion of *Coprobacter* and reduced *Lachnospiraceae_FE2018_group* (Fig. 3B) increased in the 600 mg/kg GML-treated group (*P* < 0.05). The relative abundances of *Barnesiella, Odoribacter* and *Parabacteroides* increased (*P* < 0.05) in the broilers fed with 900 mg/kg GML (Fig. 3B). In addition, except the decreased *Lachnospiraceae_FE2018_group*, 1200 mg/kg GML significantly increased the abundances of *Barnesiella, Faecalibacterium, Bacteroides, Odoribacter*, and *Parabacteroides* and *CHKC001* level in the ceca of the broilers (*P* < 0.05) relative to those in the control and broilers that received other GML doses (Fig. 3B).

**FIG 2.**
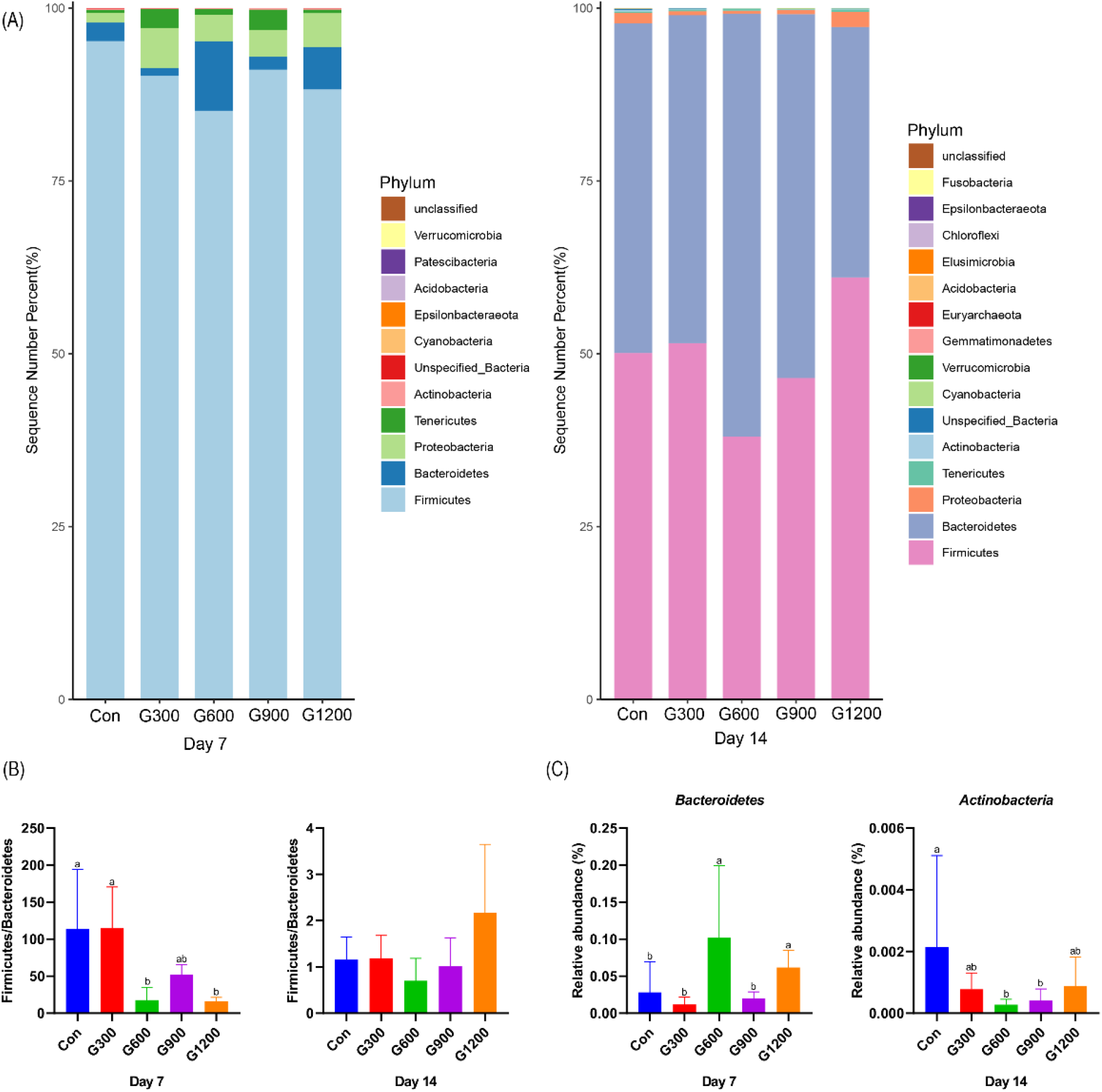
Alteration of cecal microbiota on the phylum level. (A) The top 20 phyla in the relative abundance of each group. (B) The ratio of Firmicutes/Bacteroidetes. (C) Relative abundance of *Bacteroidetes* and *Actinobacteria*. Different superscripts indicate significant differences according to the Kruskal-Wallis test (*P* < 0.05). Con, basal diet; G300, 600, 900, and 1200, basal diets complemented with 300, 600, 900, or 1200 mg/kg GML.

**FIG 3.**
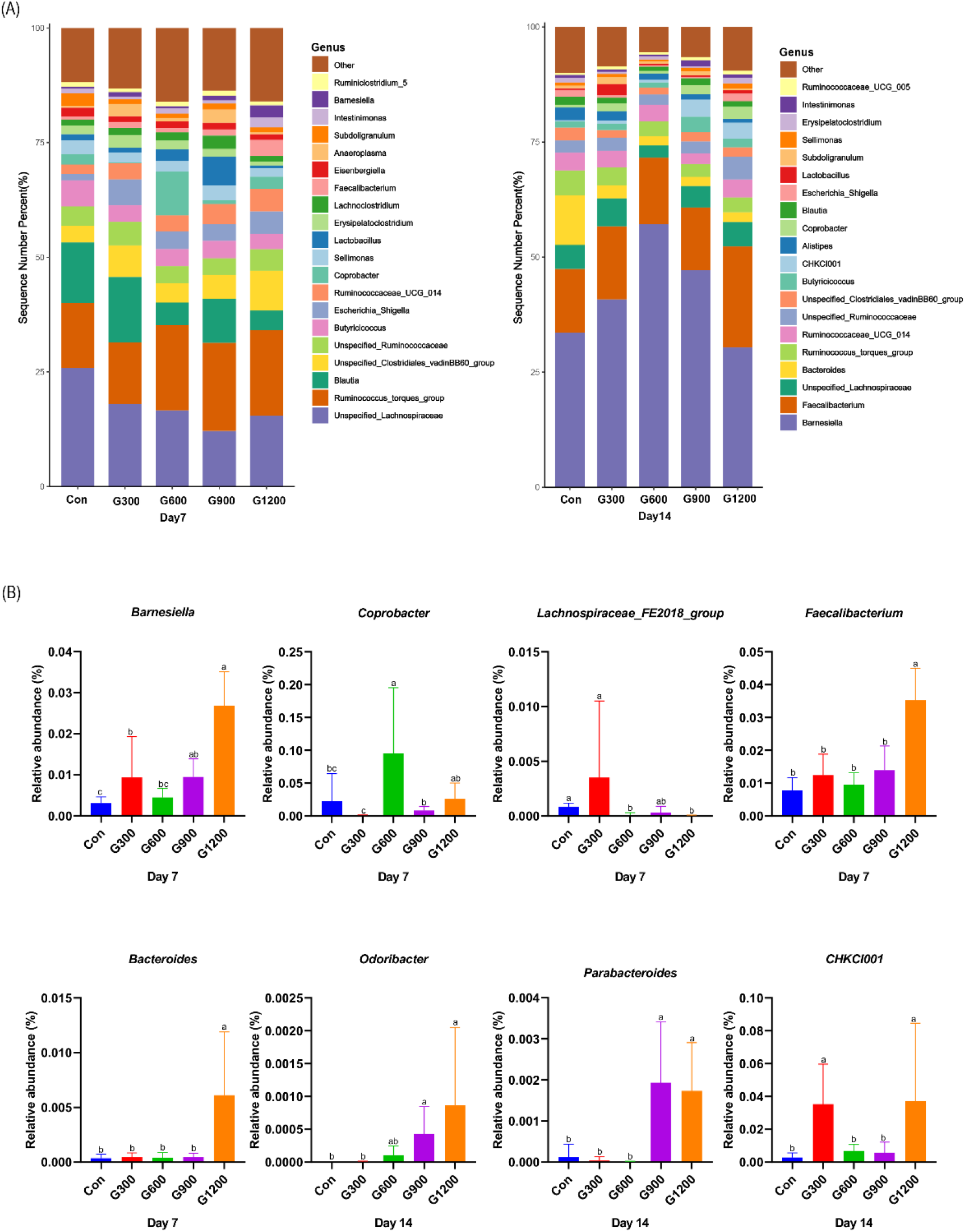
Alteration of cecal microbiota on the genus level. (A) Relative abundance of top 20 bacterial genera presents in each group. (B) The relative abundance differs significantly among groups at the genus level. Different superscripts indicate significant differences according to the Kruskal-Wallis test (*P* < 0.05). Con, basal diet; G300, 600, 900, and 1200, basal diets complemented with 300, 600, 900, or 1200 mg/kg GML.

The specific bacterial taxa associated with GML treatment was identified through linear discriminant analysis effect size (LEfSe, LDA score > 4) analysis. As shown in Fig. 4A, only the genus *Blautia* was observed to be significantly abundant in the 300 mg/kg group. The 600 mg/kg GML-treated group showed the enrichment of the genus *Coprobacter*, family *Bacteroidetes*, phylum Bacteroidia, and order Bacteroidales. The family *Lactobacillaceae* and genus *Lactobacillus* predominated in the 900 mg/kg GML group. Furthermore, the genera *Faecalibacterium, Barnesiella*, and *Intestinimonas* were enriched in the ceca of 1200 mg/kg GML-treated broilers on day 7. At the 14th day of treatment, the abundance of genus *UBA1819* increased in the 300 mg/kg GML-treated group (Fig. 4B). LEFse analysis indicated significant distinctive bacteria of family *Barnesiellaceae* and genus *Barnesiella* in the 600 mg/kg GML-treated group (Fig. 4B). Genus *Arthromitus* and *CHKCI001* were overrepresented in the 900 mg/kg group (Fig. 4B). The cecal microbiota in the 1200 mg/kg GML-fed group was characterized by phylum, family *Lachnospiraceae*, and genus *Coprobacter* (Fig. 4B). Through Spearman correlation analysis, the relationships of changes in intestinal microflora at the genus level with intestinal integrity, inflammatory factors, antioxidant enzymes, tight junction proteins, and TLR/NF-κB signal pathway were discussed (Fig. 4C). The levels of inflammatory factors were positively associated with the genera *Lachnospiraceae_UCG_008, Erysipelatoclostridium, Unspecified_Bacteria, Anaerostipes, Eubacterium_hallii_group, Eisenbergiell*a, *Lachnospiraceae_UCG_010*, and *Anaerotruncus* but negatively correlated with *Barnesiella, Faecalibacterium, Coprobacter, Odoribacter, Parabacteroides, CHKCI001*, and *Bilophila*. Intestinal integrity was possibly correlated with the genera *Coprobacter, Ruminococcaceae_UCG_010, Faecalibacterium, Barnesiella, Desulfovibrio, Bilophila, Parabacteroides, Phascolarctobacterium*, and *CHKCI001*. The abundances of *Lachnospiraceae_FE2018_group, Eubacterium_hallii_group, Ruminiclostridium_5, Enterococcus, Romboutsia*, and Caproiciproducens were negatively associated with intestinal barrier.

**FIG 4.**
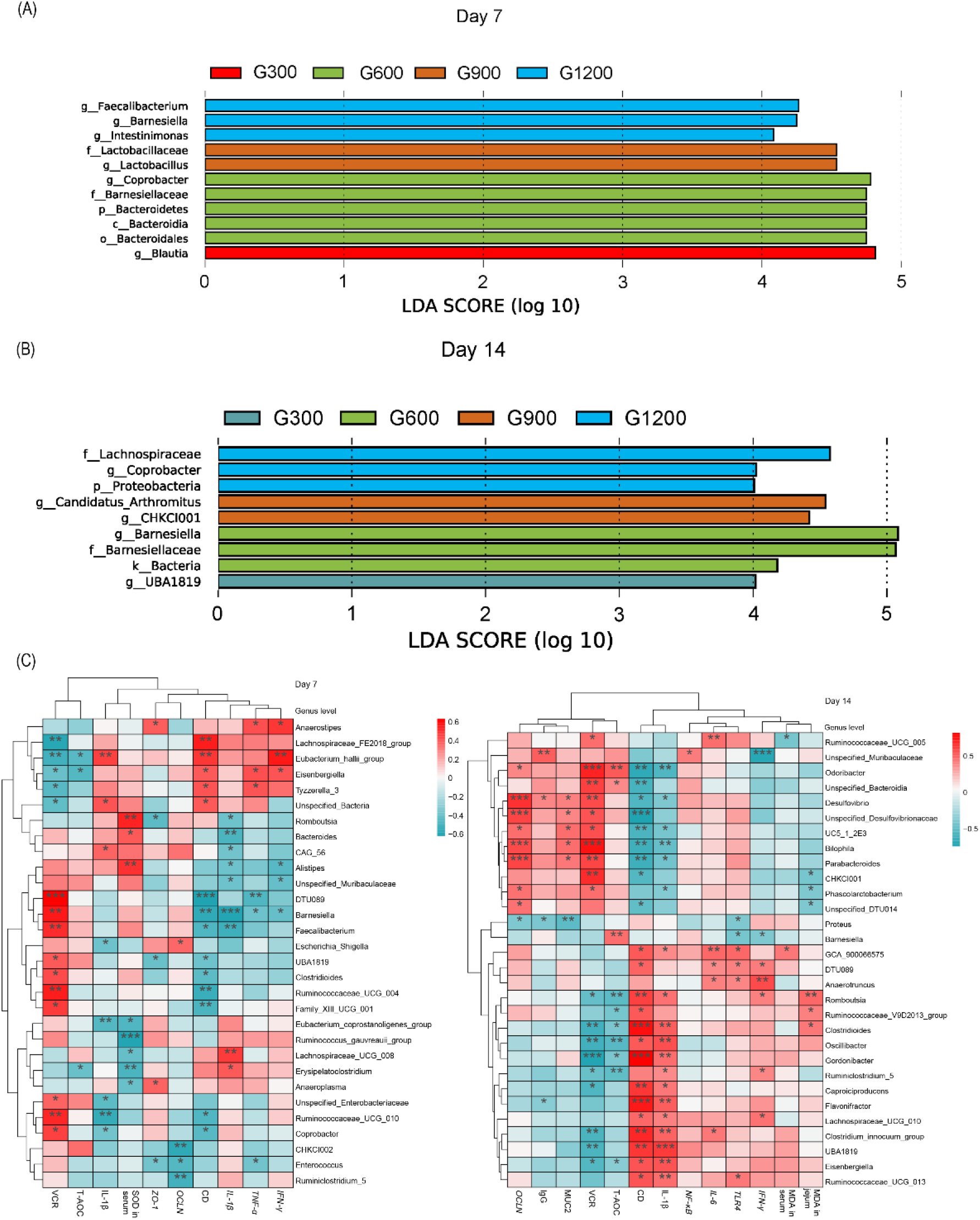
(A) LEfSe analysis of cecal microbiota (LDA score is greater than 4). (B) Heatmap of correlation among cecal microbiota, inflammatory factors, tight junctions, antioxidant enzymes, and intestinal integrity by Spearman analysis. \**P* < 0.05, \*\**P* < 0.01, \*\*\**P* < 0.001. Con, basal diet; G300, 600, 900, and 1200, basal diets complemented with 300, 600, 900, or 1200 mg/kg GML.

## Discussion

Medium-chain fatty acids (MCFA) have received widespread attention as feed additives, showing positive benefits and improving animal health, production, and feed digestibility (11, 12). Current evidence supports that MCFA and monoglycerides are generally effective in supporting growth performance and intestinal health (13). Feed intake is the basic feature that guides the growth rates of broilers (14). In the current study, dietary treatment with GML effectively increased FI rates of broiler chicks, consistent with previous findings (9, 15). A combination of gut inflammation, immune cell infiltration, and increased levels of proinflammatory cytokines often reduce FI and cause diarrhea (16). Decreased level of inflammatory factors after GML treatment may be one of the reasons for the increased FI in the present study. However, none of the supplemented treatment groups showed changes in BW, BWG, and FCR, consistent with the previous findings that GML did not alter the growth performance of broilers in the first 28 days of treatment (17). In addition, no effects were reported on the BW, BWG, and FCR after dietary supplementation with four GML levels (0, 1, 3, or 5 g/kg) during all experimental periods (3). Conversely, increased ADFI and BW and reduced FCR were observed in the 28–56-day-old yellow-feathered broilers with dietary GML (18). Broilers fed with GML had increased BWG, ADG, and feed consumption and decreased FCR (9). The positive effect of MCFA on digestibility and growth performance is the inhibition of pathogen proliferation (19).

Diets complemented with the glycerol-esters of MCFAs exert immunomodulatory effects (20). In the current study, reduced IL-1β and TNF-α and increased IgG levels were observed in the sera after dietary treatment with GML. Proinflammatory cytokines IL-6 and TNF-β levels decreased in the sera after GML supplementation, which alleviates systemic inflammatory response in high-fat-diet-fed mice (6). In addition, GML may be considered a topical anti-inflammatory agent (21). It down-regulated the gene expression of jejunal *IL-1β, IL-6*, and *IFN-γ* in the present study. Owing to the relatively stable state and long residence time in the gastrointestinal tract, an anti-inflammatory environment was induced instead of systemic inflammation after the administration of a high dose of GML (22). A similar result was observed in this study. The 1200 mg/kg GML dose showed better effects on broiler inflammation and immunity. Monoglycerides and MCFAs exhibit antimicrobial and immunomodulatory activities as additive candidates, thereby mitigating feed pathogen proliferation and improving enteric health in weaned pigs (13). An in vitro study showed that lauric acid treatment reduces the concentration of proinflammatory cytokines (TNF-α, IL-6, and IL-1β) in the culture supernatant of microglia attacked by LPS (23). T cell receptor (TCR)–induced signaling and T cell activation were suppressed by GML, which inhibits T cell-mediated cytokine storm (24). In addition, GML disrupted the lipid dynamics of human T cells, potentially reducing the TCR-induced production of cytokines. This function suggests that GML has an immunomodulatory role (5). The secretion of MIP-3 and IL-8 induced by HIV-1 was inhibited. This result supported the hypothesis that GML has an immunomodulatory effect during infection (8).

One of the central ways feed compounds affect immunity is activating NF-κB, which is an inducible central regulator of inflammatory responses involved in most innate immune receptor signaling pathways (25). In this study, GML supplementation tended to decrease jejunal *NF-κB* expression. Lauric acid and GML have minimal impacts on NF-κB activation in the absence of an LPS challenge, although a statistically significant increase can be observed at certain concentrations (25). Such increase partially explains the results of the present study. The canonical pathway of NF-κB activation involves signaling by pattern recognition receptors, such as TLR (26). Our data indicated that dietary treatment with 600 and 1200 mg/kg GML effectively reduced the relative mRNA expression of *TLR4*. Similar results were obtained in a mice experiment in which GML down-regulated *TLR2* MyD88 expression in the liver, reducing systemic inflammation in high-fat-diet fed mice (6). Lauric acid increases the activity of NF-κB through the dimerization of TLR2 and TLR1 or TLR6 and the activation of TLR4 (27).

Intestinal integrity is a key factor for preventing the invasion of pathogenic microorganisms in broilers (28). Dietary GML tends to increase the VH and VCR and decrease the CD of the jejunum, thus improving growth performance (18). In this study, dietary GML decreased CD and increased VCR in the jejuna of broilers with increasing dose. High VCR is widely regarded as a good indicator of mucosal turnover and is related to strong digestion and absorption capacity (28). VH and VCR in the duodena, jejuna, and ilea of broilers increased after 150 mg/kg GML supplementation, and this increase was considered the reason for the high metabolic rate of feed nutrients and decreased FCR (29). Natural extracts with anti-inflammatory effects restore the damaged intestinal morphology of broilers attacked by LPS (30). Therefore, the improvement in intestinal integrity can be partially explained by the decreased expression of proinflammatory cytokines and down-regulation of *TLR4/NF--κB* signal transduction. After GML was given to the mice, the normal expression levels of *ZO-1, occludin, claudin-1, jam-1*, and *muc 2* suggested that GML maintained the mucosal barrier and intestinal health (22). In this study, dietary GML effectively benefited the jejunal muc 2 content and up-regulated *ZO-1* and *occludin* expression, demonstrating the beneficial effect of GML on intestinal barrier of broilers. The TNF-α and IFN-γ are related to the reduction in epithelial barrier function and increases the permeability of the mucosal barrier (31). The reduction in serum TNF-α level and decrease in jejunal *IFN-γ* expression in this study supported the idea that GML mediates intestinal barrier function through a mechanism associated with the attenuation of intestinal inflammation.

Oxidative stress is an important factor for the destruction of mucosal barrier function (32). In the current study, reduced MDA content and increased T-SOD and T-AOC activity in the sera and jejuna indicated that GML reduced lipid peroxidation and improved the antioxidant capacity of the broilers. This result is in a line with a previous study (33). In laying hens, increase in SOD level and reduced glutathione and MDA levels were observed, suggesting that GML can decrease lipid peroxidation and enhance the antioxidant capacities of broilers. These features are beneficial to growth performance (17). The peroxidation level in the meat of GML-fed chickens was reduced and proportional to the increase in dietary additive concentration (9). Significant connections between inflammation and oxidative stress have been found. These processes induce each other reciprocally, thereby establishing a vicious cycle that perpetuates and propagates inflammatory response (34). In addition, the proinflammatory signaling cascades triggered by TLR engagement enhance the expression of iNOS, implying that TLR activation may result in oxidative stress (35). Therefore, the improvement in antioxidant capacity in the GML-treated broilers may be related to lowered inflammatory response and down-regulated TLR4/NF--κB pathway. Reduced oxidative stress was beneficial for relieving inflammation.

Intestinal microbiota contributes to the maintenance of intestinal physiological structure and function, which are considered relevant to intestinal inflammation, barrier function, and growth performance of a host (36). Our results showed that dietary GML modulated the microbial composition in the ceca of the broilers. A rich community of species enhances the stability of the intestinal microecology and may be related to reduced sensitivity to bacterial invasion and intestinal inflammation (37). Therefore, the alleviated intestinal inflammation of broilers may be associated with the modulated structure and increased diversity of intestinal microbial community structure after GML addition. At the phylum level, GML mainly altered the relative frequency of *Bacteroidetes* and *Actinobacteria* in the ceca of the broilers. Increase in *Actinobacteria* and *Firmicutes*/*Bacteroidetes* ratio is the pattern of an impaired intestinal barrier (38). In this study, the cecal microbiota in GML-treated broilers was characterized by decreased level of *Actinobacteria* and *Firmicutes*/*Bacteroidetes* ratio, which may be the potential reasons for the improved intestinal barrier. In addition, 1200 mg/kg GML increased the amount of cecal *Bacteroidetes*, which mainly contributes to the fermentation of indigestible carbohydrates to butyrate (39) and exerts beneficial effects on mucosal barrier integrity through its anti-inflammatory effects (40). At the genus level, the relative abundance of *Barnesiella, Coprobacter, Lachnospiraceae, Faecalibacterium, Bacteroides, Odoriacter, Parabacteroides*, and *CHKCI001* were higher in the GML-treated groups, especially under the dose of 1200 mg/kg. Mice fed with 400 and 800 mg/kg GML had higher abundances of *Barnesiella*, which was considered to be positively associated with a healthy state (22). *Lachnospiraceae* can promote health by producing host nutrients and providing an energy supply to the colonic epithelium, as well as maintaining host immune homeostasis (41). *Parabacteroides* are associated with T-cell differentiation by enhancing and maintaining the IL-10-producing Treg cells (42). *Barnesiella, Faecalibacterium, Bacteroides, Odoribacter*, and *Parabacteroides* were positively correlated with the level of SCFAs and produced butyrate, which exerted immunomodulatory and anti-inflammatory effects by mediating the homeostasis of colonic regulatory T cell populations and promoted intestinal integrity (43). In addition, butyrate with antioxidant properties modulates inflammatory response by inhibiting NF-κB and provides energy to the intestinal epithelial cells (38). A significant impact was reported on host health and physiology after GML supplementation, which directly acts on the intestinal microbiota and considerably affects metabolism and immunity (44). Therefore, the increased abundance of butyrate-producing bacteria after GML addition was associated with alleviated intestinal inflammation and improved intestinal morphology of the broilers.

In conclusion, dietary GML ameliorated intestinal morphology and barrier function of broiler chicks by ameliorating inflammation and promoting antioxidant status. Our results confirmed the immunomodulatory, antioxidant, and anti-inflammatory properties of GML, which may be associated with the suppression of the TLR4/NF-κB signaling pathway. The altered structure of the cecal microbiota manipulated by GML may be the main reason for the promotion of intestinal health in the broilers.

## Material and methods

### Animals, experimental design and management

All experimental procedures were approved by the Ethics Committee of the Shandong Agricultural University and carried out according to the Guidelines for Experimental Animals of the Ministry of Science and Technology (Beijing, People’s Republic of China). All feeding and euthanasia procedures were performed with full consideration of animal welfare. A total of 360 one-day-old broilers (Arbor Acres) with an average weight of 45.7 g were randomly divided into five groups as follows: basal diet (control) and basal diets supplemented with 300, 600, 900, or 1200 mg/kg GML, which was purchased from Henan Zhengtong Food Technology Co., Ltd (Henan, China) with purity of more than 90%. The additive dosage of GML was optimized according to previous studies (6, 45, 46). Each group contained six replicates (cage) of 12 broilers per cage. The ingredients and nutrients levels in the basal diet were formulated according to standards of the National Research Council 2012 (Table 1). All broilers were weighed and randomly assigned to 30 metal cages (70 cm × 70 cm × 40 cm), which were equipped with feeders and nipple drinkers. Broilers with similar initial weights were reared in an environmentally controlled room. The temperature was 35 °C initially, and then gradually decreased to 28 °C until the end of the 14-day experiment period. In the first 3 days, average relative humidity was maintained at approximately 70%, and thereafter maintained between 55% and 65%. In the first week, the broilers were kept under 23 h of light and 1 h of darkness, which were then gradually reduced to 20 and 4 h, respectively.

### Growth performance

On days 7 and 14, feed consumption in each replicate and body weight were recorded. Body weight gain (BWG) and feed intake (FI) were calculated subsequently. Spilled feed was carefully collected and weighed for the correction of the final FI data. Feed conversion rate (FCR) was defined as FI:BWG. The data of mortality were recorded and included in the FCR calculation.

### Sampling

Two broilers per replicate were randomly selected for sampling after growth performance was determined on 7 and 14 days of age. Blood samples were collected from wing veins and negotiated to glass tubes without anticoagulants, then centrifuged at 3000 rpm for 10 min at 4 °C. Serum was obtained and stored at −20 °C for biochemical analysis. Broilers were slaughtered by cervical dislocation after blood samples were obtained. Approximately 2 cm segments were excised from the jejunum (from the entry point of the bile duct to the Meckel’s diverticulum), flushed repeatedly with cold saline solution, and immediately immersed in 4% paraformaldehyde solution for histological examination. Tissue samples (1–2 g) were collected from the jejunum, rapidly frozen in liquid nitrogen, and stored at −80 °C for molecular analysis. The cecum was collected on ice, frozen quickly in liquid nitrogen, transported to the laboratory in a dry-ice bag, and then stored at −80 °C for further microbial analysis.

### Jejunal morphology analysis

Jejunum segments were fixed in 4% paraformaldehyde solution for 24 h, dehydrated, and embedded in paraffin. Tissue sections with 5 μm thickness were cut using a microtome (Leica RM2235, Leica Biosystems Inc., Buffalo Grove, USA), fixed on slides, and stained with hematoxylin and eosin. The images of the jejunum were analyzed with ImageJ analysis software (Version 1.47, Bethesda, MD, USA). Ten intact villi were selected randomly from each section for morphology measurement. Villus height (VH) was gauged from the tip of the villus to the villus–crypt junction. Crypt depth (CD) was defined as the depth of the invagination between adjacent villi. Villus height-to-crypt depth ratio (VCR) was calculated. The mean value of ten values attributed to individual broilers was used in statistical analysis.

### Biochemical assay of serum and jejunum

Immune response status in the sera was estimated by detecting the levels of interleukin 1 beta (IL-1β), interleukin 6 (IL-6), tumor necrosis factor-alpha (TNF-α), and immunoglobulin G (IgG) with ELISA kits (MLBIO Co., Ltd., Shanghai, China). All determination procedures were performed strictly according to the manufacturer’s instructions. The inter- and intra-assay coefficients of variation (CVs) were less than 10%. The jejunal samples were weighed accurately (0.3 g), homogenized with 2.7 ml phosphate-buffered saline in a weight (g):volume (ml) ratio of 1: 9. The homogenates were centrifuged at 1000 *g* for 10 min at 4 °C, and the supernatants were collected for the detection of secreted immunoglobulin A (sIgA) and mucin 2 levels with ELISA kits (MLBIO Co., Ltd., Shanghai, China). The results were expressed as pg/mg of protein.

### Antioxidant assay of serum and jejunum

Malondialdehyde (MDA) levels, total antioxidant capacity (T-AOC), and total superoxide dismutase (T-SOD) activity were measured in each serum and jejunal homogenate with diagnostic kits (intra-assay CV < 5%; inter-assay CV < 8%) purchased from Nanjing Jiancheng Biotechnology Institute (Nanjing, China) according to the manufacturer’s instructions. The results were normalized to protein concentration in each jejunal homogenate.

### RNA isolation and real-time quantitative PCR

The total RNA of the jejunum was isolated using Trizol reagent (Invitrogen Biotechnology Inc., CA, USA). RNA quality was evaluated through 1% agarose gel electrophoresis. The reverse transcription of 1 μg of total RNA was performed using PrimeScript® RT reagent kit with gDNA Eraser (RR047A, Takara Bio Inc., Dalian, China). The gene expression was determined through RT-PCR. TB Green Premix Ex Taq (RR820A, Takara Bio Inc., Dalian, China) and ABI 7500 real-time PCR systems (Applied Biosystems, CA, USA) were used. The reaction program was as follows: predenaturation at 95 °C for 10 s, then denaturation at 95 °C for 5 s for a total of 40 cycles, and finally annealing and extension at 60 °C for 40 s. Each reaction was repeated three times, and the primer sequences are shown in Table 2. The amplification efficiency of the primers was calculated with a standard curve. The specificity of the amplified products was verified with the melting curve. The relative expression of the target gene was analyzed through the 2^−ΔΔCt^ method after normalization against the geometric mean of the expression of β-actin, glyceraldehyde-3-phosphate dehydrogenase (GAPDH), and TATA-binding protein (TBP).

### 16S rRNA sequencing and analysis

Total DNA was extracted from cecal contents with an E.Z.N.A.® Soil DNA kit (Omega Bio-Tek, Norcross, GA, U.S.) according to the manufacturer’s instructions. DNA purity and concentration were evaluated with a Nano Drop2000 spectrophotometer (Thermo Scientific, Wilmington, USA), and DNA integrity was detected through 1% agarose gel electrophoresis. Bacterial 16S rRNA gene spanning the V3-V4 hypervariable regions were amplified with primers 338F (5′-ACTCCTACGGGAGGCAGCAG-3′) and 806R (5′-GGACTACHVGGGTWTCTAAT-3′) with a PCR system. PCR reactions were performed in triplicate with a 20 μl mixture consisting of 4 μl of 5 × FastPfu Buffer, 2 μl of 2.5 mM dNTPs, 0.8 μl of each primer (5 μM), 0.4 μl of FastPfu polymerase, and 10 ng of template DNA. The amplification programs were set in ABI GeneAmp® 9700 system (ABI, USA) as follows: 3 min at 95 °C, 27 cycles of 30 s at 95 °C, 55 °C for 30 s, and 72 °C for 45 s, and 72 °C for 10 min. The PCR products were detected through 2% agarose gel electrophoresis and purified with an AxyPrep DNA gel recovery kit (Axygen Biosciences, Union City, CA, USA) and then quantified with a QuantiFluor(tm)-ST blue fluorescence quantitative system (Promega, USA). Purified amplicons were pooled in equimolar amounts, and their paired-end reads were sequenced on an Illumina MiSeq PE300 platform (Illumina, San Diego, USA) by Majorbio Bio-Pharm Technology Co., Ltd. (Shanghai, China). Raw fastq files obtained through MiSeq sequencing were demultiplexed, quality-filtered, trimmed, de-noised with trimmomatic, and merged according to the overlapping relationship by FLASH (version 1.2.11, https://ccb.jhu.edu/software/FLASH/index.shtml). The filtered reads were clustered into operational taxonomic units (OTUs) with a 97 % sequence identity with UPARSE (version 7.1, http://www.drive5.com/uparse/). Chimera was removed during clustering. The OTU representative sequence was analyzed with the RDP Classifier (version 2.2, http://sourceforge.net/projects/rdp-classifier/) against the Silva 16S rRNA database (release119, http://www.arb-silva.de) at a confidence threshold of 70%.

### Statistical analysis

Data were presented as mean ± SD. All data were checked for normality with Shapiro–Wilk test (95% confidence level). Statistical differences between groups were analyzed with one-way ANOVA with Tukey’s multiple comparisons or by the non-parametric factorial Kruskal-Wallis test. SPSS software (SPSS 26.0, SPSS, Chicago, USA) was used. Differences were considered significantly different at *P* < 0.05. Probability 0.05 < *P* < 0.1 were defined as tendencies.

## ACKNOWLEDGMENT

This work was supported by the Natural Science Foundation of Shandong Province (ZR2020MC170), the National Key R&D Program of China (2018YFD0501401-3), and the Shandong Province Agricultural Industry Technology (SDAIT-11-08).

## REFERENCE

1. Sánchez-Cordón PJ, Montoya M, Reis AL, Dixon LK. 2018. African swine fever: A re-emerging viral disease threatening the global pig industry. Veterinary journal (London, England : 1997) 233:41–48.

2. Kogut MH. 2017. Issues and consequences of using nutrition to modulate the avian immune response. Journal of Applied Poultry Research 26:605–612.

3. Amer SA, A-Nasser A, Al-Khalaifah HS, AlSadek DMM, Abdel Fattah DM, Roushdy EM, Sherief WRIA, Farag MFM, Altohamy DE, Abdel-Wareth AAA, Metwally AE. 2020. Effect of Dietary Medium-Chain α-Monoglycerides on the Growth Performance, Intestinal Histomorphology, Amino Acid Digestibility, and Broiler Chickens’ Blood Biochemical Parameters. Animals : an open access journal from MDPI 11.

4. Wang C, Shao C, Fang Y, Wang J, Dong N, Shan A. 2021. Binding loop of sunflower trypsin inhibitor 1 serves as a design motif for proteolysis-resistant antimicrobial peptides. Acta Biomater 124:254–269.

5. Zhang MS, Sandouk A, Houtman JCD. 2016. Glycerol Monolaurate (GML) inhibits human T cell signaling and function by disrupting lipid dynamics. Scientific reports 6:30225.

6. Zhao M, Jiang Z, Cai H, Li Y, Mo Q, Deng L, Zhong H, Liu T, Zhang H, Kang JX, Feng F. 2020. Modulation of the Gut Microbiota during High-Dose Glycerol Monolaurate-Mediated Amelioration of Obesity in Mice Fed a High-Fat Diet. mBio 11.

7. Fortuoso BF, Galli GM, Griss LG, Armanini EH, Silva AD, Fracasso M, Mostardeiro V, Morsch VM, Lopes LQS, Santos RCV, Gris A, Mendes RE, Boiago MM, Da Silva AS. 2020. Effects of glycerol monolaurate on growth and physiology of chicks consuming diet containing fumonisin. Microb Pathog 147:104261.

8. Welch JL, Xiang J, Okeoma CM, Schlievert PM, Stapleton JT. 2020. Glycerol Monolaurate, an Analogue to a Factor Secreted by, Is Virucidal against Enveloped Viruses, Including HIV-1. mBio 11.

9. Fortuoso BF, Dos Reis JH, Gebert RR, Barreta M, Griss LG, Casagrande RA, de Cristo TG, Santiani F, Campigotto G, Rampazzo L, Stefani LM, Boiago MM, Lopes LQ, Santos RCV, Baldissera MD, Zanette RA, Tomasi T, Da Silva AS. 2019. Glycerol monolaurate in the diet of broiler chickens replacing conventional antimicrobials: Impact on health, performance and meat quality. Microb Pathog 129:161–167.

10. Pan D, Yu Z. 2014. Intestinal microbiome of poultry and its interaction with host and diet. Gut microbes 5:108–119.

11. Jackman JA, Hakobyan A, Zakaryan H, Elrod CC. 2020. Inhibition of African swine fever virus in liquid and feed by medium-chain fatty acids and glycerol monolaurate. Journal of animal science and biotechnology 11:114.

12. Baltic B, Starčević M, Djordjevic J, Mrdović B, Markovic R. 2017. Importance of medium chain fatty acids in animal nutrition. IOP Conference Series: Earth and Environmental Science 85:012048.

13. Jackman JA, Boyd RD, Elrod CC. 2020. Medium-chain fatty acids and monoglycerides as feed additives for pig production: towards gut health improvement and feed pathogen mitigation. Journal of animal science and biotechnology 11:44.

14. Abdollahi MR, Zaefarian F, Ravindran V. 2018. Feed intake response of broilers: Impact of feed processing. Animal Feed Science and Technology 237:154–165.

15. Mustafa N. 2018. Biochemical Trails Associated with Different Doses of Alpha-Monolaurin in Chicks. Advances in Animal and Veterinary Sciences 7.

16. Pié S, Lallès JP, Blazy F, Laffitte J, Sève B, Oswald IP. 2004. Weaning is associated with an upregulation of expression of inflammatory cytokines in the intestine of piglets. The Journal of nutrition 134:641–647.

17. Liu T, Li C, Zhong H, Feng F. 2021. Dietary medium-chain α-monoglycerides increase BW, feed intake, and carcass yield in broilers with muscle composition alteration. Poultry science 100:186–195.

18. Liu T, Tang J, Feng F. 2020. Glycerol monolaurate improves performance, intestinal development, and muscle amino acids in yellow-feathered broilers via manipulating gut microbiota. Appl Microbiol Biotechnol 104:10279–10291.

19. Hong SM, Hwang JH, Kim IH. 2012. Effect of Medium-chain Triglyceride (MCT) on Growth Performance, Nutrient Digestibility, Blood Characteristics in Weanling Pigs. Asian-Australasian journal of animal sciences 25:1003–1008.

20. Masmeijer C, Rogge T, van Leenen K, De Cremer L, Deprez P, Cox E, Devriendt B, Pardon B. 2020. Effects of glycerol-esters of saturated short-and medium chain fatty acids on immune, health and growth variables in veal calves. Preventive veterinary medicine 178:104983.

21. Peterson ML, Schlievert PM. 2006. Glycerol monolaurate inhibits the effects of Gram-positive select agents on eukaryotic cells. Biochemistry 45:2387–2397.

22. Mo Q, Fu A, Deng L, Zhao M, Li Y, Zhang H, Feng F. 2019. High-dose Glycerol Monolaurate Up-Regulated Beneficial Indigenous Microbiota without Inducing Metabolic Dysfunction and Systemic Inflammation: New Insights into Its Antimicrobial Potential. Nutrients 11.

23. Nishimura Y, Moriyama M, Kawabe K, Satoh H, Takano K, Azuma Y-T, Nakamura Y. 2018. Lauric Acid Alleviates Neuroinflammatory Responses by Activated Microglia: Involvement of the GPR40-Dependent Pathway. Neurochemical research 43:1723–1735.

24. Zhang MS, Tran PM, Wolff AJ, Tremblay MM, Fosdick MG, Houtman JCD. 2018. Glycerol monolaurate induces filopodia formation by disrupting the association between LAT and SLP-76 microclusters. Science signaling 11.

25. Sivinski SE, Mamedova LK, Rusk RA, Elrod CC, Swartz TH, McGill JM, Bradford BJ. 2020. Development of an macrophage screening system on the immunomodulating effects of feed components. Journal of animal science and biotechnology 11:89.

26. Sun S-C. 2017. The non-canonical NF-κB pathway in immunity and inflammation. Nature reviews Immunology 17:545–558.

27. Lee JY, Zhao L, Youn HS, Weatherill AR, Tapping R, Feng L, Lee WH, Fitzgerald KA, Hwang DH. 2004. Saturated fatty acid activates but polyunsaturated fatty acid inhibits Toll-like receptor 2 dimerized with Toll-like receptor 6 or 1. The Journal of biological chemistry 279:16971–16979.

28. Jiang J, Qi L, Lv Z, Jin S, Wei X, Shi F. 2019. Dietary Stevioside Supplementation Alleviates Lipopolysaccharide-Induced Intestinal Mucosal Damage through Anti-Inflammatory and Antioxidant Effects in Broiler Chickens. Antioxidants (Basel, Switzerland) 8.

29. Liu M, Chen X, Zhao H, Feng F. 2018. Effect of Dietary Supplementation with Glycerol Monolaurate on Growth Performance, Digestive Ability and Chicken Nutritional Components of Broilers. Food Science.

30. Shang Y, Regassa A, Kim JH, Kim WK. 2015. The effect of dietary fructooligosaccharide supplementation on growth performance, intestinal morphology, and immune responses in broiler chickens challenged with Salmonella Enteritidis lipopolysaccharides. Poultry science 94:2887–2897.

31. Cromarty R, Archary D. 2020. Inflammation, HIV, and Immune Quiescence: Leveraging on Immunomodulatory Products to Reduce HIV Susceptibility. AIDS research and treatment 2020:8672850.

32. S L, Chaudhary S, R S R. 2017. Hydroalcoholic extract of Stevia rebaudiana bert. leaves and stevioside ameliorates lipopolysaccharide induced acute liver injury in rats. Biomedicine & pharmacotherapy = Biomedecine & pharmacotherapie 95:1040–1050.

33. Valentini J, Da Silva AS, Fortuoso BF, Reis JH, Gebert RR, Griss LG, Boiago MM, Lopes LQS, Santos RCV, Wagner R, Tavernari FC. 2020. Chemical composition, lipid peroxidation, and fatty acid profile in meat of broilers fed with glycerol monolaurate additive. Food chemistry 330:127187.

34. Lugrin J, Rosenblatt-Velin N, Parapanov R, Liaudet L. 2014. The role of oxidative stress during inflammatory processes. Biol Chem 395:203–230.

35. Lucas K, Maes M. 2013. Role of the Toll Like receptor (TLR) radical cycle in chronic inflammation: possible treatments targeting the TLR4 pathway. Mol Neurobiol 48:190–204.

36. Wang W, Li Z, Han Q, Guo Y, Zhang B, D’Inca R. 2016. Dietary live yeast and mannan-oligosaccharide supplementation attenuate intestinal inflammation and barrier dysfunction induced by Escherichia coli in broilers. The British journal of nutrition 116:1878–1888.

37. Ott SJ, Musfeldt M, Wenderoth DF, Hampe J, Brant O, Fölsch UR, Timmis KN, Schreiber S. 2004. Reduction in diversity of the colonic mucosa associated bacterial microflora in patients with active inflammatory bowel disease. Gut 53:685–693.

38. Olejniczak-Staruch I, Ciążyńska M, Sobolewska-Sztychny D, Narbutt J, Skibińska M, Lesiak A. 2021. Alterations of the Skin and Gut Microbiome in Psoriasis and Psoriatic Arthritis. International journal of molecular sciences 22.

39. Islam MR, Lepp D, Godfrey DV, Orban S, Ross K, Delaquis P, Diarra MS. 2019. Effects of wild blueberry (Vaccinium angustifolium) pomace feeding on gut microbiota and blood metabolites in free-range pastured broiler chickens. Poultry science 98:3739–3755.

40. Yu M, Mu C, Zhang C, Yang Y, Su Y, Zhu W. 2018. Marked Response in Microbial Community and Metabolism in the Ileum and Cecum of Suckling Piglets After Early Antibiotics Exposure. Frontiers in microbiology 9:1166.

41. Shi S, Qi Z, Jiang W, Quan S, Sheng T, Tu J, Shao Y, Qi K. 2020. Effects of probiotics on cecal microbiome profile altered by duck Escherichia coli 17 infection in Cherry Valley ducks. Microb Pathog 138:103849.

42. Arpaia N, Campbell C, Fan X, Dikiy S, van der Veeken J, deRoos P, Liu H, Cross JR, Pfeffer K, Coffer PJ, Rudensky AY. 2013. Metabolites produced by commensal bacteria promote peripheral regulatory T-cell generation. Nature 504:451–455.

43. Zhou Y, Wang Y, Quan M, Zhao H, Jia J. 2021. Gut Microbiota Changes and Their Correlation with Cognitive and Neuropsychiatric Symptoms in Alzheimer’s Disease. Journal of Alzheimer’s disease : JAD doi:10.3233/JAD-201497.

44. Jiang Z, Zhao M, Zhang H, Li Y, Liu M, Feng F. 2018. Antimicrobial Emulsifier-Glycerol Monolaurate Induces Metabolic Syndrome, Gut Microbiota Dysbiosis, and Systemic Low-Grade Inflammation in Low-Fat Diet Fed Mice. Molecular nutrition & food research 62.

45. Amer SA, A AN, Al-Khalaifah HS, AlSadek DMM, Abdel Fattah DM, Roushdy EM, Sherief W, Farag MFM, Altohamy DE, Abdel-Wareth AAA, Metwally AE. 2020. Effect of Dietary Medium-Chain α-Monoglycerides on the Growth Performance, Intestinal Histomorphology, Amino Acid Digestibility, and Broiler Chickens’ Blood Biochemical Parameters. Animals (Basel) 11.

46. Ren C, Wang Y, Lin X, Song H, Zhou Q, Xu W, Shi K, Chen J, Song J, Chen F, Zhang S, Guan W. 2020. A Combination of Formic Acid and Monolaurin Attenuates Enterotoxigenic Escherichia coli Induced Intestinal Inflammation in Piglets by Inhibiting the NF-κB/MAPK Pathways with Modulation of Gut Microbiota. J Agric Food Chem 68:4155–4165.

